# Glucokinase activity suppresses hepatic cholesterol synthesis and triglyceride accumulation: A new model for the effects of the GKRP P466L common human variant

**DOI:** 10.64898/2026.04.07.717049

**Authors:** Dominic Santoleri, Sarah Traynor, Ryan Calhoun, Matthew J. Gavin, David Merrick, Patrick Seale, Paul M. Titchenell

**Affiliations:** Institute of Diabetes, Obesity, and Metabolism, Perelman School of Medicine, University of Pennsylvania; Biochemisry, Biophysics, and Chemical Biology Graduate Group, Perelman School of Medicine, University of Pennsylvania; Department of Medicine, Perelman School of Medicine, University of Pennsylvania; Department of Cell and Developmental Biology, Perelman School of Medicine, University of Pennsylvania; Department of Physiology, Perelman School of Medicine, University of Pennsylvania

**Keywords:** Glucokinase (GCK), Glucokinase Regulatory Protein (GKRP), GKRP P446L, cholesterol synthesis, fatty liver

## Abstract

**Objective:** Glucokinase Regulatory Protein (GKRP) controls the activity of Glucokinase (GCK) to regulate liver glucose uptake and storage. Coding variants in *GCKR*, the gene encoding GKRP, strongly associate with fatty liver disease, hypertriglyceridemia, and hypercholesterolemia. Here, we sought to investigate the mechanisms by which a common GKRP variant affects hepatic lipid and cholesterol metabolism.

**Methods:** We developed mouse models to examine how the human GKRP P446L variant influences liver and systemic metabolism. Endogenous Gckr expression was ablated in adult mouse hepatocytes, together with re-expression of either human GKRP P446L or the reference GKRP protein. We assessed body weight, adiposity, systemic glucose homeostasis, and hepatic metabolites in mice expressing reference GKRP or GKRP P446L under multiple metabolic conditions. To determine whether the effects of GKRP P446L may result from reduced GCK activity, we analyzed mice with liver-specific deletion of *Gck*.

**Results:** Hepatic expression of GKRP P446L resulted in reduced GKRP and GCK protein levels and elevated serum cholesterol. Hepatic deletion of *Gck* in mice recapitulated several effects of GKRP P446L, including increased hepatic cholesterol and triglyceride content. The elevated cholesterol was associated with increased cholesterogenic gene expression and cholesterol synthesis. Hepatic expression of an alternative hexokinase (HKII) normalized the effects of GCK-deficiency, suggesting that impaired glucose phosphorylation underlies the phenotype.

**Conclusions:** The GKRP P446L variant reduced GKRP protein abundance, and diminished GCK activity while increasing cholesterol levels. Loss of GCK elevated cholesterol and hepatic triglyceride levels. Collectively, these findings demonstrate that GCK suppresses hepatic cholesterol synthesis and lipid accumulation, suggesting that reduced GCK activity underlies the metabolic abnormalities associated with the GKRP P446L variant.

**Highlights:**

- The GKRP P446L variant reduces GKRP protein abundance and diminishes GCK activity.
- Expression of GKRP P446L in mouse hepatocytes increases serum cholesterol levels.
- GCK activity suppresses cholesterogenic gene expression and cholesterol synthesis.

## 1. Introduction

Glucokinase (GCK) plays a central role in hepatic glucose metabolism by catalyzing the conversion of glucose to glucose-6-phosphate (G6P), thereby directing glucose towards metabolic utilization or storage as glycogen [1,2]. *GCKR* encodes for Glucokinase Regulatory Protein (GKRP), which binds and inhibits GCK in hepatocytes by sequestering it in the nucleus [3,4]. When hepatic glucose levels are low, such as during fasting, GCK adopts a low-activity, super-open conformation that is bound and stabilized by GKRP [5–7]. As glucose levels rise, such as after a meal, glucose promotes a conformational shift of GCK to a closed, active state, releasing it from GKRP and enabling glucose phosphorylation and uptake [8,9]. Thus, GKRP coordinates the transition between glucose uptake and glucose production depending on glucose availability while maintaining a reserve pool of GCK that can rapidly respond to changing metabolic conditions [10].

Genome wide association studies (GWAS) have identified variants in *GCKR* that strongly associate with decreased blood glucose, elevated serum triacylglycerides (TAG) and cholesterol [11–15]. Moreover, these variants correlate with an increased risk of fatty liver disease but protection against Type 2 Diabetes. In particular, the single nucleotide polymorphism (SNP), rs1260326 [c.1403 C>T, p.P446L] is a common *GCKR* variant [16], with minor allele frequencies ranging between 0.1 and 0.4 in different populations [16,17], strongly correlated to the above conditions.

*In vitro* studies on the *GCKR* SNP rs126036 (henceforward referred to as GKRP P446L) demonstrate that the variant has reduced binding affinity for GCK [18–20]. These findings gave rise to the prevailing model that the P446L variant inappropriately increases hepatic GCK activity, promoting glucose uptake and *de novo* lipogenesis, thereby elevating triglyceride and cholesterol levels. Studies of pharmacological glucokinase activators (GKAs) support this model, as they improve glycemic control in hyperglycemic or diabetic patients but frequently cause hepatic steatosis and hypertriglyceridemia [21,22]. However, these effects may be partly attributable to GCK activation in pancreatic β-cells, where it serves as the glucose sensor controlling insulin secretion. Timing GKA treatment to feeding periods also reduces steatosis, suggesting that inappropriate activation of GCK during fasting contributes more to the increase in steatosis than generalized overactivation of the pathway [23].

Recently, the Agius lab generated a germline knock-in mouse model expressing the murine equivalent of the human GKRP P446L variant [24]. These mice exhibit reduced postprandial blood glucose and insulin levels, along with elevated serum cholesterol and hepatic TAG content. Interestingly, the P446L variant caused a marked loss of GKRP protein in both mouse and human livers accompanied by reduced GCK expression and activity, resembling phenotypes observed in GKRP knockout models [25,26].

We investigated the effects of the GKRP P446L variant by generating adult-onset, liver-specific *Gckr* knockout mice and re-expressing either the reference human GKRP protein or the GKRP P446L variant using hepatocyte-specific AAV vectors. Expression of GKRP P446L resulted in reduced GKRP and GCK protein levels, accompanied by glucose intolerance and elevated serum cholesterol. Notably, loss of GCK was sufficient to elevate hepatic TAG and cholesterol levels. The increase in cholesterol coincided with enhanced *de novo* cholesterol synthesis. Collectively, these findings suggest that reduced, rather than increased, GCK activity drives the hepatic steatosis and hypercholesterolemia associated with the GKRP P446L variant in humans.

## 2. Results

### 2.1. Loss of GKRP in hepatocytes decreases GCK levels and activity

The GKRP P446L variant associates with elevated circulating lipids and cholesterol and reduced risk of insulin resistance (**Fig. S1**). To determine how the GKRP P446L variant affects hepatic metabolism, we generated mice where we could delete the endogenous *Gckr* gene and re-express either the reference human GKRP or the GKRP P446L variant specifically in hepatocytes. We created *Gckr* floxed mice by inserting *loxP* sites flanking exon 5 of *Gckr*, the deletion of which produces a frameshift and premature termination codon (**Fig. S2**). Injecting *Gckr*^*fl/fl*^ mice with hepatocyte-specific AAV-TBG-Cre ablated *Gckr* mRNA expression and GKRP protein in the liver (L-GKRP-KO) compared with mice injected with control AAV-TBG-GFP (**Fig. 1A** and **1B**). Notably, GCK protein levels also demonstrated reduced expression in the livers of L-GKRP-KO mice, particularly in the refed state following a 6-hour fast (**Fig. 1B**). This finding is consistent with prior work indicating that loss of GKRP impairs the feeding-induced restoration of GCK protein expression [25,26]. Liver *Gck* mRNA levels did not change between refed L-GKRP-KO and control mice, supporting the role of GKRP in stabilizing GCK protein (**Fig. 1B**). Loss of GKRP also reduced the expression of *Mlxipl* and *Pklr* (**Fig. 1B**), genes whose transcription depends on glucose phosphorylation [27,28].

**Figure 1:**
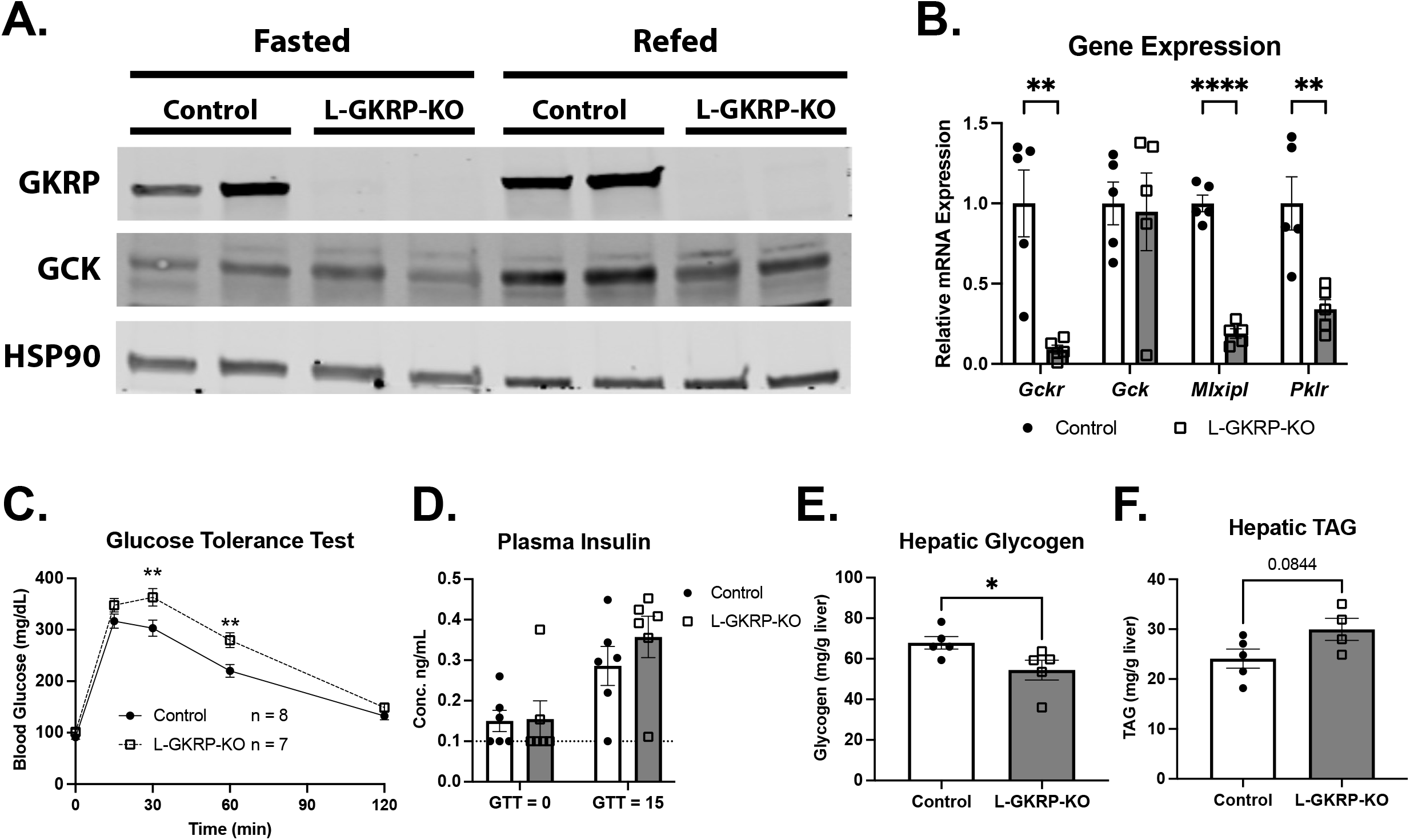
Hepatic GKRP is required for normal liver glucokinase (GCK) activity and glucose utilization. **A)** Western blot analysis of GKRP and GCK in livers from control and L-GKRP-KO mice under fasting (overnight) or refeeding conditions (overnight fast followed by 6 hours of refeeding). HSP90 was used as a loading control. **B)** mRNA levels of *Gckr, Gck, Mlxipl*, and *Pklr* in livers from refed control and L-GKRP-KO mice after 1 week on HCD (n = 5 per group). **C)** Glucose tolerance test performed on control and L-GKRP-KO mice after 3 days on HCD (n = 8/9 control/KO). **D)** Plasma insulin levels of mice from (C) at baseline (fasted) and 15 minutes after glucose injection (n = 6 per group). Points highlighted in red measured below the limit of detection (0.1 ng/mL) of the insulin ELISA and were thus set to 0.1 ng/mL. **E**,**F)** Hepatic glycogen (E) and TAG (F) levels in control and L-GKRP-KO mice under fasting (overnight) or refeeding conditions (overnight fast followed by 6 hours of refeeding) after 1 week on HCD (n = 5 per group). Data are represented as mean ± SEM. Statistical significance was determined using Student’s t test (B, E, and F) or 2-way ANOVA with either Šidák *post hoc* test (C) or Fisher’s Least Significant Difference (D). * p < 0.05, ** p < 0.01, *** p < 0.001, **** p < 0.0001.

To functionally evaluate GCK activity, we measured glucose tolerance in control and L-GKRP-KO mice fed a high-carbohydrate diet (HCD) for 3 days and measured glucose tolerance because while GKRP deletion reduces GCK expression on normal chow, it does not alter glucose homeostasis unless nutritionally challenged [25]. L-GKRP-KO mice exhibited impaired glucose clearance (**Fig. 1C**) without a significant change in plasma insulin levels (**Fig. 1D**). Consistent with reduced hepatic glucose clearance and utilization, livers from overnight-fasted and 6-hour-refed L-GKRP-KO mice after 1 week on HCD exhibited reduced glycogen content compared with controls (**Fig. 1E**). Hepatic triacylglycerol (TAG) levels also trended modestly upward in L-GKRP-KO mice after 1 week on HCD (**Fig. 1F**). Collectively, these findings demonstrate that normal GCK expression and activation in response to refeeding requires hepatic GKRP.

### 2.2 GKRP P446L reduces hepatic GCK expression, disrupts glucose homeostasis, and elevates cholesterol

We next assessed the effects of the GKRP P446L variant by injecting *Gckr*^*fl/fl*^ mice with AAV-TBG-Cre together with AAV-TBG vectors expressing either the human reference GKRP protein (hGKRP) or the P446L variant. Beginning two weeks after injection, the mice consumed a high-fat diet (HFD) for 13 weeks. Strikingly, the P446L variant showed markedly reduced protein expression compared with hGKRP, accompanied by substantially lower GCK protein levels (**Fig. 2A**). P446L-expressing mice modestly gained more body weight and displayed a trend for an increase in fat mass compared with hGKRP-expressing mice (**Fig. 2B** and **2C**). Liver mass normalized to body weight did not differ significantly between groups (**Fig. 2D**). After 9 weeks of HFD feeding, P446L-expressing mice displayed impaired glucose tolerance compared with hGKRP-expressing mice (**Fig. 2E**) with no change in insulin levels at baseline or 15 minutes after glucose administration (**Fig. 2F**). Hepatic glycogen content also trended lower in P446L mice (**Fig. 2G**). Collectively, these results indicate that the P446L variant destabilizes GKRP, resulting in reduced GCK protein levels and activity.

**Figure 2:**
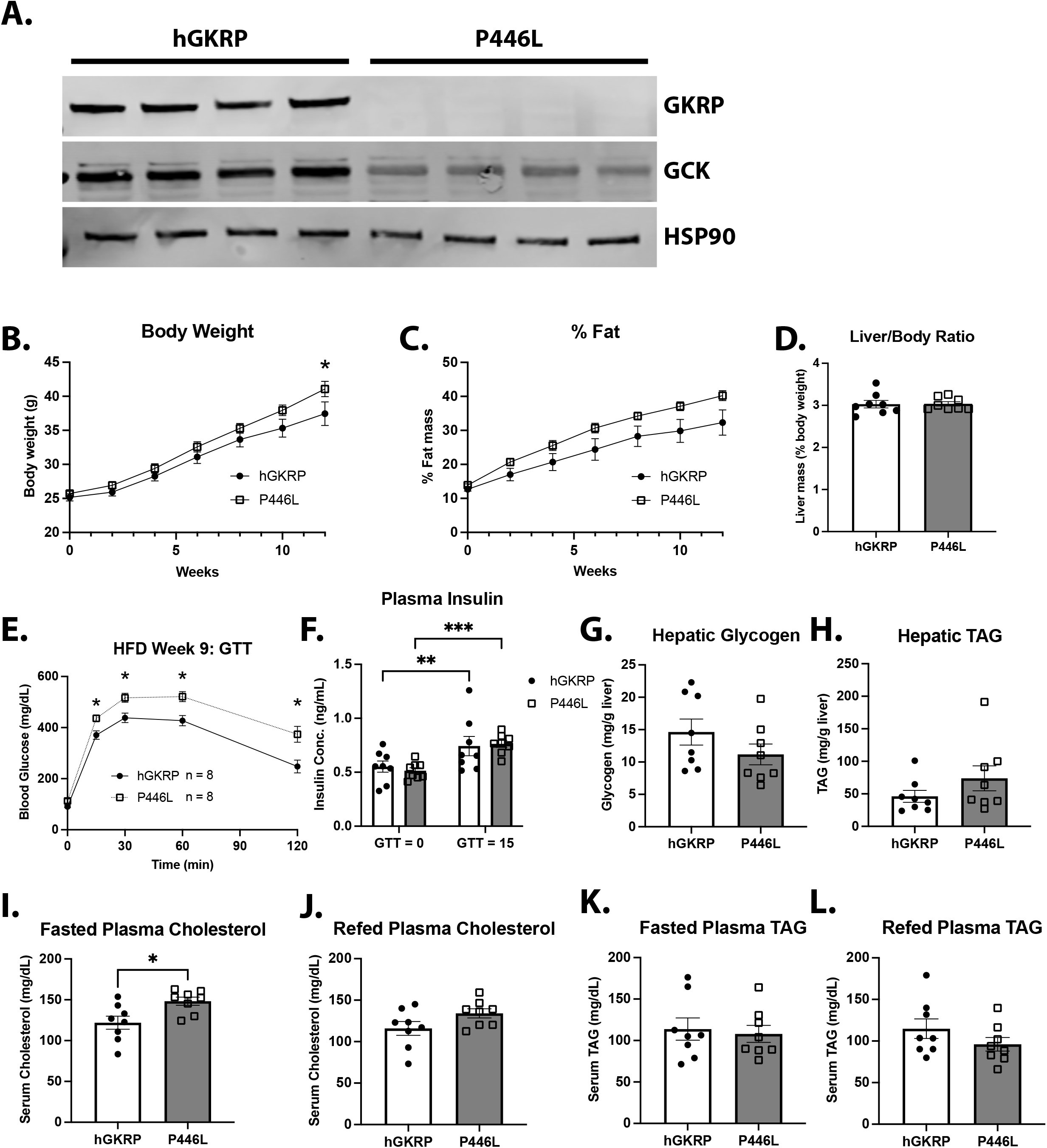
The GKRP-P446L variant impairs glucose homeostasis and increases plasma cholesterol. **A)** Western blot analysis of GKRP and GCK in livers from L-GKRP-KO mice expressing either human GKRP (hGKRP) or the P446L variant. Samples were collected from mice after an overnight fast followed by 4 hours of refeeding after 13 weeks of HFD feeding. HSP90 was used as a loading control. **B-D)** Body weight (B), percent fat mass (C), and liver mass (% body weight) of hGKRP- and P446L-expressing mice over 13 weeks of HFD feeding (n = 8 per group). **E)** Glucose tolerance test on hGKRP- and P446L-expressing mice after 9 weeks on HFD diet (n = 8 per group). **F)** Plasma insulin levels in mice from (E) at baseline (fasted) and 15 minutes after glucose injection. Samples that measured below the assay’s limit of detection (0.1 ng/mL) were set to 0.1 (marked with a dashed line on the graph). **G-L)** L-GKRP-KO mice expressing hGKRP and hGKRP-P446L were fed a HFD for 13 weeks (n = 8 for both groups) **G)** Hepatic glycogen content after an overnight fast and refeeding for 4 hours. **H)** Hepatic TAG after overnight fast and refeeding for 4 hours. **I)** Plasma cholesterol after an overnight fast. **J)** Plasma cholesterol after an overnight fast and refeeding for 4 hours. **K)** Plasma TAG from overnight fasted mice. **L)** Plasma TAG after overnight fast and refeeding for 4 hours. Data are represented as mean ± SEM. Statistical significance was determined using either 2-way ANOVA with either Šidák *post hoc* test (B, C, and E) or Fisher’s Least Significant Difference (F) or Student’s t test (D, G-L). * p < 0.05, ** p < 0.01, *** p < 0.001.

Additionally, we observed a trend toward increased hepatic TAG levels and significantly elevated fasting serum cholesterol in P446L mice compared with hGKRP-expressing mice (**Fig. 2H** and **2I**). Plasma cholesterol levels in P446L mice normalized after 4 hours of refeeding (**Fig. 2J**), indicating that feeding can reverse the fasting hypercholesterolemia. Plasma TAG levels did not differ between groups under either overnight-fasted or 4-hour-refed conditions (**Fig. 2K** and **3L**). These findings are similar to those reported in germline P446L knock-in mice, which also exhibited increased serum cholesterol and hepatic TAG levels without changes in circulating TAG [24]. Overall, these data demonstrate that the lipid and cholesterol effects of the P446L variant in mice associate with reduced GCK activity.

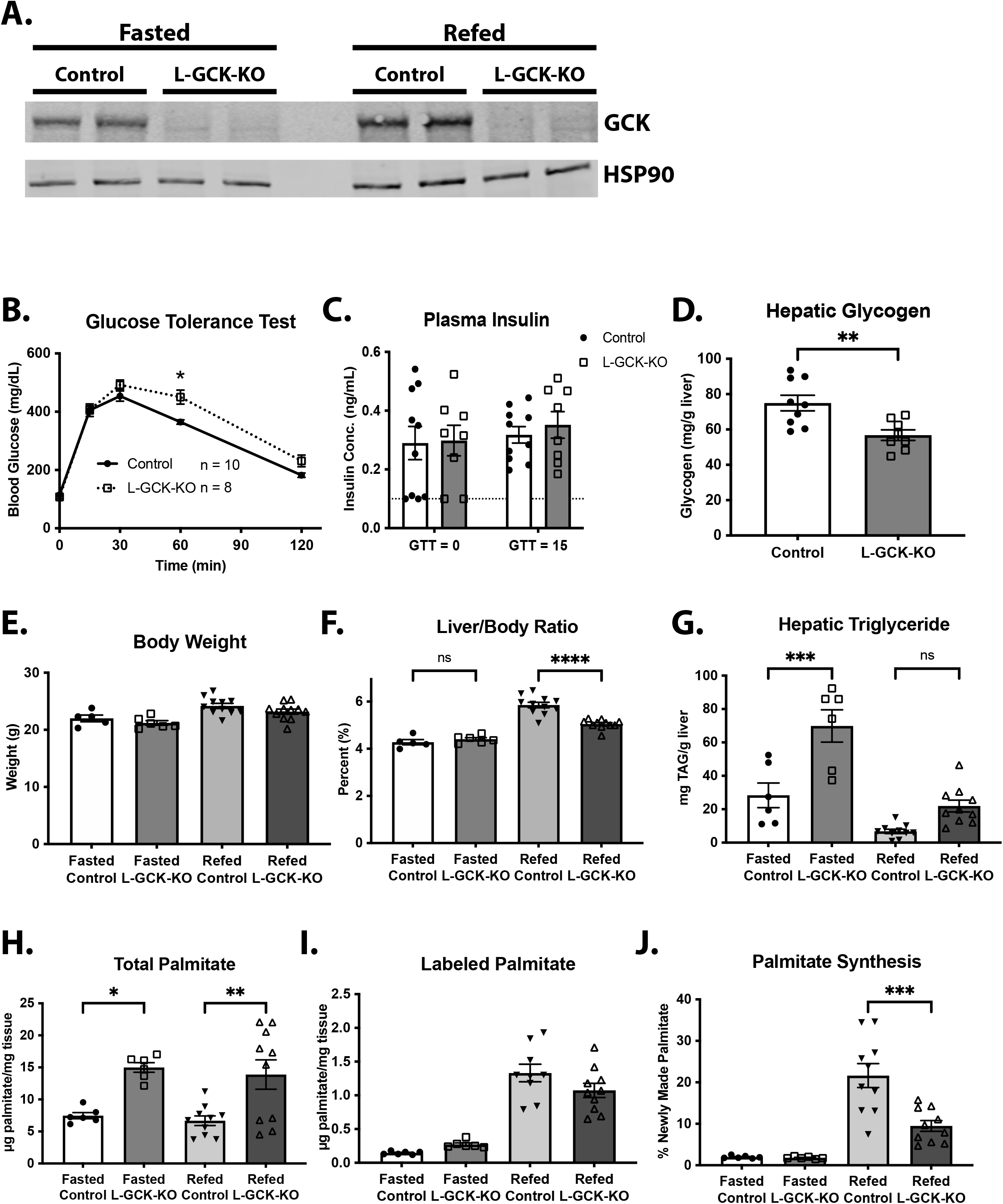
Error! No text of specified style in document.: Glucokinase suppresses hepatic TAG accumulation. **A)** Western blot analysis of GCK in livers from control and L-GCK-KO mice fed a HCD for 1 week. Samples were obtained from mice under fasting (overnight) or refeeding conditions (overnight fast followed by 4 hours of refeeding). HSP90 was used as a loading control. **B)** Glucose tolerance test on control and L-GCK-KO mice after 3 days on HCD (n = 10/8 control/KO). **C)** Plasma insulin levels in mice from (B) at baseline (fasted) and 15 minutes after glucose injection (n = 10/8 control/KO). Samples that measured below the assay’s limit of detection (0.1 ng/mL) were set to 0.1 (marked with a dashed line on the graph). **D)** Hepatic glycogen content in control and L-GCK-KO mice under refeeding conditions (n = 9/8 control/KO). **E-J)** Livers were obtained from control and L-GCK-KO mice under fasting (overnight) or refeeding conditions (overnight fast followed by 6 hours of refeeding) after HCD for 1 week. (control fasted n = 6, L-Gck-KO fasted n = 6, control refed n = 10, L-GCK-KO refed n = 8). **E)** Body weight. **F)** Liver mass (% of body weight). **G)** Hepatic TAG content. **H)** Total hepatic palmitate. **I)** Total D_2_O labeled hepatic palmitate. **J)** % labeled palmitate (of total hepatic palmitate). Data are represented as mean ± SEM. Statistical significance was determined using 2-way ANOVA with either Šidák *post hoc* test (B) or Fisher’s Least Significant Difference (C), Student’s t test (D), or 1-way ANOVA with Tukey’s *post hoc* test (E-J). * p < 0.05, ** p < 0.01, *** p < 0.001, **** p < 0.0001.

### 2.3. GCK suppresses hepatic triglyceride accumulation and cholesterol synthesis

To test whether a reduction of GCK activity *per se* mediates the effects of hGKRP P446L expression on hepatic metabolism, we deleted *Gck* specifically in the liver. We injected *Gck*^*fl/fl*^ mice with AAV-TBG-Cre to ablate GCK expression (L-GCK-KO) or control AAV (AAV-TBG-GFP) and fed them a high-carbohydrate diet (HCD). We collected livers after either an overnight fast or an overnight fast followed by 6 hours of refeeding. All these mice received intraperitoneal D_2_O to monitor hepatic *de novo* lipogenesis (DNL) via quantifying deuterium incorporation into palmitate. Western blot analysis showed efficient deletion of GCK in L-GCK-KO livers (**Fig. 3A**). After 3 days of HCD, L-GCK-KO mice exhibited impaired glucose tolerance compared with control mice (**Fig. 3B**) without changes in plasma insulin levels at baseline or 15 minutes after glucose administration (**Fig. 3C**). Additionally, L-GCK-KO mice had significantly reduced hepatic glycogen content (**Fig. 3D**). After 1 week of HCD, body weight did not differ between control and L-GCK-KO mice under either fasted or refed conditions (**Fig. 3E**). However, refed L-GCK-KO mice had significantly lower liver mass relative to body weight (**Fig. 3F**). Hepatic TAG levels increased (∼3-fold) in L-GCK-KO mice compared with controls under fasting conditions and showed a trend toward higher levels in the refed state (**Fig. 3G**). Further, we observed an increase in the levels of palmitate in L-GCK-KO livers under both fasted and refed conditions (**Fig. 3H**). Despite this increase, both groups demonstrated comparable deuterium incorporation into palmitate, indicating reduced fractional DNL rates in L-GCK-KO mice (**Fig. 3I** and **3J**).

To assess how loss of GCK affects hepatic gene programming, we performed RNA sequencing (RNA-seq) on liver samples from L-GCK-KO and control mice after 6 hours of refeeding following an overnight fast (**Fig. 4A**). Gene ontology analysis of differentially expressed genes (fold change ≥ 2 or ≤ −2; adjusted p value < 0.05) identified genes involved in sterol and cholesterol biosynthesis as the most strongly altered pathways in L-GCK-KO livers while downregulated genes did not strongly fit into any pathway (**Fig. 4B** and **Fig. S3**). Genes encoding many key enzymes in the cholesterol synthesis pathway, including *Pcsk9, Sqle, Fdft1, Fdps*, and *Hmgcr*, were robustly upregulated in L-GCK-KO livers relative to control (**Fig. 4A**). We validated the induction of these genes by qRT-PCR analysis of livers from a larger cohort of refed L-GCK-KO and control mice (**Fig. 4C**).

**Figure 4:**
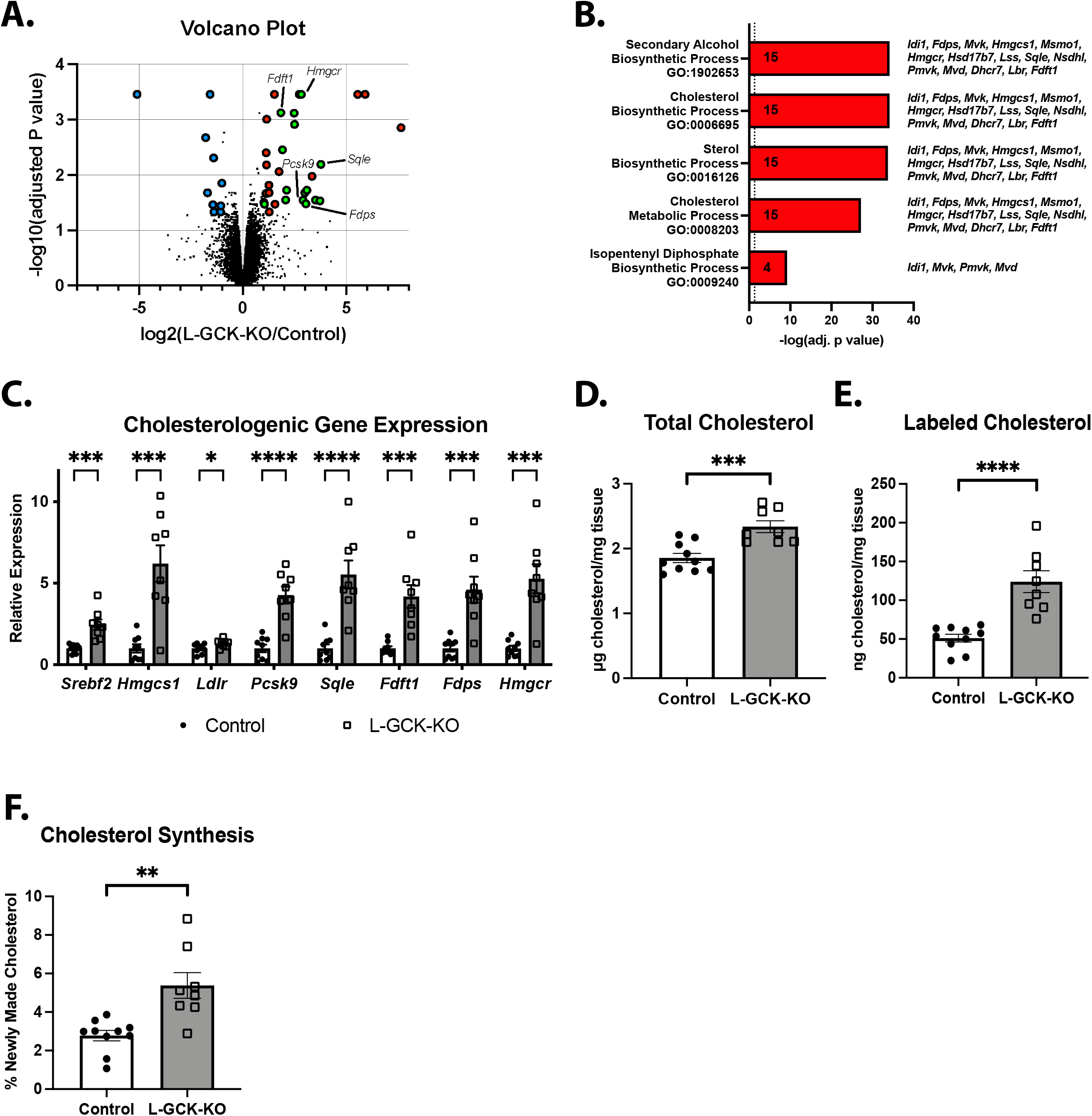
Glucokinase is required to suppress hepatic cholesterol synthesis. **A)** Volcano plot showing differentially expressed genes identified from RNA-sequencing analysis of livers from control and L-GCK-KO mice fed a HCD for 1 week. Samples were obtained from mice that were fasted overnight followed by refeeding for 6 hours. (n = 3/4 control/KO). Red labels: genes upregulated in KO vs. control (FC ≥ 2, adjusted p value < 0.05, total = 44). Green labels: upregulated cholesterol synthesis genes (total = 16). Blue labels: genes downregulated in KO vs. control (FC ≤ -2, adjusted p value < 0.05, total = 15). **B)** Gene ontology analysis of significantly up regulated genes (GO Biological Process 2026). Numbers in each bar represent the number of differentially regulated genes and the list of those genes is presented next to it. **C)** mRNA levels of cholesterogenic genes in livers from control and L-GCK-KO mice fed a HCD for 1 week (n = 9/8 control/KO). **D-F)** Control and L-GCK-KO mice were fed a HCD for 1 week. Samples were obtained from mice that were fasted overnight followed by refeeding for 24 hours. (n=10/8 control/KO). **D)** Total hepatic cholesterol. **E)** D_2_O-labeled hepatic cholesterol. **F)** % of labeled cholesterol (of total cholesterol). Data are represented as mean ± SEM. Statistical significance was determined using Student’s t test (**C-F**). * p < 0.05, ** p < 0.01, *** p < 0.001, **** p < 0.0001.

To determine whether these transcriptional changes translate into increased cholesterol synthesis, we measured *de novo* synthesis using deuterium oxide (D_2_O) labeling. Mice received a D_2_O injection and had *ad libitum* access to food for 24 hours while drinking 4% D_2_O-enriched water. L-GCK-KO mice displayed elevated total hepatic cholesterol levels (**Fig. 4D**) together with about a 2.5-fold increase in deuterium incorporation into cholesterol (**Fig. 4E**), reflecting a marked increase in cholesterol synthesis (**Fig. 4F**). Together, these results demonstrate that loss of GCK drives hepatic cholesterol synthesis, providing a potential explanation for the elevated cholesterol levels observed in humans carrying the GKRP P446L variant.

### 2.4. Glucose phosphorylation suppresses hepatic cholesterol synthesis

Recent work suggests that GCK also functions in the nucleus to regulate gene transcription [29]. To determine whether glucose phosphorylation suppresses cholesterol synthesis and TAG accumulation, we generated an AAV vector to express hexokinase II (HKII), a distinct hexokinase isoform, specifically in hepatocytes (AAV-TBG-HKII). Because HKII catalyzes glucose phosphorylation but does not engage GCK-specific regulatory interactions, such as binding to GKRP [30], we expected its expression in hepatocytes to restore glucose uptake without engaging in GCK-specific protein interactions.

Injection of L-GCK-KO mice with AAV-TBG-HKII resulted in robust HKII protein expression in liver, a tissue that does not normally express HKII (**Fig. 5A**). HKII expression normalized glucose tolerance (**Fig. 5B** and **5C**) and prevented the reduction in hepatic glycogen content observed in L-GCK-KO mice (**Fig. 5D**), demonstrating that HKII restored hepatic glucose utilization. Moreover, HKII partially restored the expression of *Mlxipl* and *Pklr*, two glucose-6-phosphate-regulated genes with reduced expression in L-GCK-KO livers (**Fig. 5E**). Strikingly, HKII expression reduced the elevated expression of cholesterol synthesis genes (e.g., *Srebf2, Hmgcr, Fdps, Fdft1, Sqle*, and *Pcsk9*) in L-GCK-KO livers and suppressed the associated increase in *de novo* cholesterol synthesis (**Fig. 5F-I**). Taken together, these results strongly suggest that the glucose-phosphorylating activity of GCK suppresses cholesterol synthesis.

**Figure 5:**
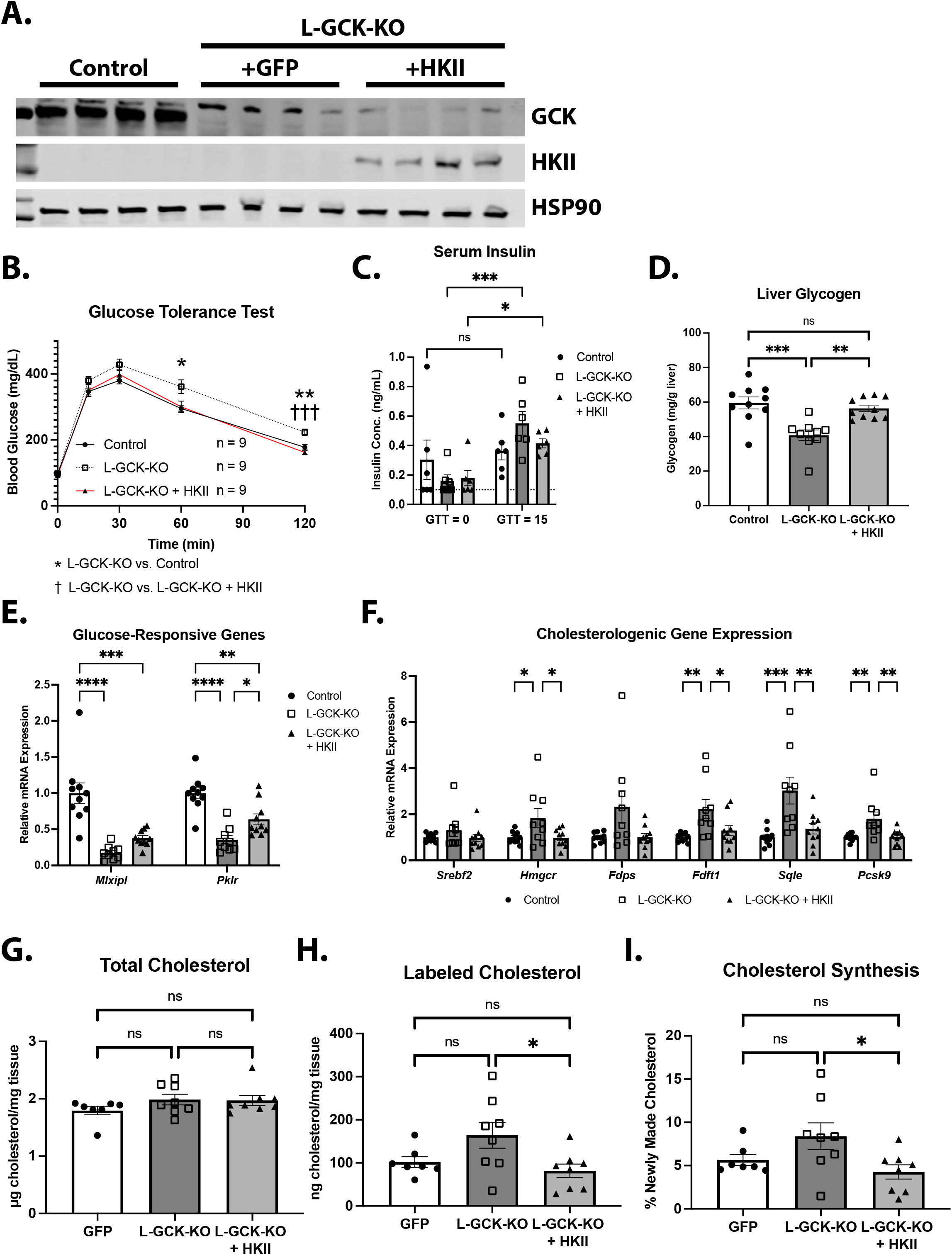
HKII overexpression normalizes cholesterol synthesis in L-GCK-KO mice. **A)** Western blot analysis of GCK and HKII in livers from control, L-GCK-KO, and HKII-overexpressing L-GCK-KO mice after 1 week on HCD. Samples were obtained from mice that were fasted overnight followed by refeeding for 24 hours. HSP90 was used as a loading control. **B)** Glucose tolerance test on control, L-GCK-KO, and HKII-overexpressing L-GCK-KO mice after 3 days on HCD (n = 9 per group). **C)** Plasma insulin levels from mice in (B) at baseline (fasting) and 15 minutes after glucose infusion (n = 9 for all groups). Samples that measured below the assay’s limit of detection (0.1 ng/mL) were set to 0.1 (marked with a dashed line on the graph). **D-I)** Control, L-GCK-KO, and HKII-overexpressing L-GCK-KO mice fed a HCD for 1 week. Measurements were performed on mice that were fasted overnight, followed by refeeding for 24 hours. **D)** Hepatic glycogen content. **E)** Liver mRNA levels of *Mlxipl* and *Pklr*. **F)** Liver mRNA levels of cholesterogenic genes. (D-F control n = 10, L-GCK-KO n = 9, +HKII n = 10) G) Total hepatic cholesterol. H) D_2_O-labeled hepatic cholesterol. I) % of labeled cholesterol (of total cholesterol). (G-I control n = 7, L-GCK-KO n = 8, +HKII n = 8). Data are represented as mean ± SEM. Statistical significance was determined using either 2-way ANOVA with Tukey’s *post hoc* test (B-C) or 1-way ANOVA with Tukey’s *post hoc* test (D-I). * p < 0.05, ** p < 0.01, *** p < 0.001, **** p < 0.0001.

## 3. Discussion

In this study, we developed and employed mouse models to investigate the physiological consequences of the GKRP P446L variant linked to elevated plasma triglyceride and cholesterol levels. Our results reveal that GKRP P446L is unstable, leading to reduced levels of functional GKRP protein and consequently diminished expression and activity of its partner enzyme GCK. Reduced GCK activity, in turn, leads to increased hepatic TAG and cholesterol levels. Together, these data are inconsistent with the canonical view that GKRP P446L drives excess lipid production due to a cell intrinsic increase in GCK activity. Instead, they support a new mechanistic model in which the GKRP P446L variant disrupts hepatic lipid homeostasis by destabilizing GCK and reducing its enzymatic activity.

Previous research has linked hepatic glucose metabolism to the suppression of cholesterol synthesis. Zhang, *et al*. demonstrated that the G6P-regulated transcription factor, carbohydrate response element binding protein β (ChREBP-β) suppresses cholesterol synthesis by promoting ubiquitination and degradation of the cholesterogenic transcription factor SREBP2 [31]. Consistent with this framework, both our L-GKRP-KO and L-GCK-KO models exhibited reduced hepatic expression levels of *Mlxipl*, the gene encoding ChREBP-β. These findings suggest that GCK may suppress cholesterol synthesis by generating G6P, which activates ChREBP-β-dependent pathways. Supporting this model, expression of HKII in GCK-deficient liver was sufficient to normalize cholesterogenic gene levels and cholesterol levels. Because HKII catalyzes glucose phosphorylation, but does not share the regulatory interactions of GCK, these results suggest that the suppression of cholesterol synthesis is mediated downstream of glucose phosphorylation. Although our experiments cannot fully exclude the possibility that GCK exerts additional non-catalytic effects, such as regulating other proteins through direct interaction, this possibility appears unlikely given the substantial structural differences between HKII and GCK.

Our findings are consistent with a recent study by Ford *et al*., which similarly reported that constitutive expression of the GKRP P446L variant in mice leads to reduced hepatic GKRP and GCK protein levels [24]. In both studies, the effects of the P446L variant on hepatic cholesterol synthesis were more pronounced than the changes in TAGs. While our data largely concur with this prior paper regarding the effects on lipids, there are still some discrepancies between models. Most notably, Ford *et al*. saw no effect of the P446L variant on glucose tolerance after 8 weeks on HFD. There are several differences between the models that can explain these variations. Chiefly, our model is an inducible, liver-specific overexpression system whereas the model presented by Ford *et al*. is a congenital, whole-body knock-in. Our model would not account for developmental effects or compensatory mechanisms that may appear in a congenital model. Additionally, the high fat diets used in either experiment differ in nutritional content (60% fat here compared to 45%), which could alter metabolic homeostasis. Alternatively, our model is hepatocyte specific and these observed differences to the whole-body model could indicate metabolic roles of GKRP in other tissues that should be investigated. GKRP expression occurs almost exclusively in hepatocytes, but low-level expression has been reported in other tissues, such as the pancreas [18].

The effects of the P446L variant on cholesterol levels in mice contrasts with human genetic studies, where the P446L variant is most strongly associated with elevated circulating TAG levels. The basis for this difference is unclear but may reflect species-specific differences in lipid metabolism or the influence of additional genetic and environmental modifiers present in human populations. For example, humans and mice utilize distinct mechanisms to transport lipids, particularly cholesterol [32,33]. Humans shuttle cholesterol from the liver in low- and very-low-density lipoproteins (LDL and VLDL) using the cholesteryl ester transfer protein (CETP). In contrast, mice lack CETP and primarily rely on high-density lipoproteins (HDL) for cholesterol transport [32]. Because VLDL particles carry substantially more TAG than HDL, an increased cholesterol burden in humans may promote VLDL secretion and elevate circulating TAG, whereas mice may not respond similarly.

While deletion of GCK recapitulated many of the lipid phenotypes observed in P446L-expressing mice, the knockout model did not fully mirror the P446L phenotype. One explanation for this difference is that the P446L model results in partial loss of GCK activity, whereas the L-GCK-KO model completely abolishes GCK function. This distinction may account for the differences in metabolic phenotype between the models. Notably, the P446L mice exhibited the most pronounced increases in hepatic triglyceride and cholesterol levels during fasting, whereas L-GCK-KO mice displayed elevated lipid levels even in the fed state. Feeding and insulin signaling are known to increase GCK expression in hepatocytes [34–36]. Thus, although GCK is destabilized in the P446L model, feeding-induced increases in its expression may partially restore GCK activity. In the absence of GKRP-mediated inhibition, even modest amounts of active GCK could generate sufficient G6P to partially suppress hepatic cholesterol synthesis during feeding. Consistent with this, hepatic glycogen levels were not reduced in P446L mice.

In summary, our findings demonstrate that the GKRP P446L variant results in reduced GKRP protein abundance and diminished GCK activity in the liver, leading to increased cholesterol synthesis and dysregulated lipid metabolism. These results provide a mechanistic explanation for how a common human genetic variant influences hepatic lipid homeostasis. Further elucidation of the pathways through which GCK-dependent glucose metabolism restrains cholesterol synthesis may reveal new therapeutic opportunities to reduce dyslipidemia and susceptibility to liver disease.

## 4. Methods

### 4.1. Mice

All mouse experiments were reviewed and approved by the University of Pennsylvania IACUC in accordance with the guidelines of the NIH. All mice in this study were male and maintained on the C57BL/6J background. The *Gck* flox mice were described previously [36]. The *Gckr* flox mice were created using CRISPR-assisted gene targeting. 20 bp gRNA sequences were designed flanking *Gckr* exon 5 using the crRNA design tool on the Integrated DNA Technologies (IDT) website (5’GGCGGACTCCAAACAAGTTA 3’ and 5’TTAGTCTCCAACATCCCTTG 3’). Each crRNA along with the Alt-R™ CRISPR-Cas9 tracrRNA (IDT #1072533) was resuspended in injection buffer (10 mM Tris-HCl, 0.1 mM EDTA, pH 7.5) to a concentration of 1 µg/µL. Each crRNA was mixed with tracrRNA in equimolar ratios and annealed per manufacturer’s instructions to create gRNA. Single stranded DNA repair templates were also created with IDT containing similar sequences to the target regions and loxP sites as well as an additional homology region (∼500 bp) on either end of each template. The loxP sites were placed within the gRNA target sequences/PAM sites to disrupt further Cas9 cutting once the homology template was incorporated into the genome. A 100 µL injection mixture containing two gRNAs (final concentration 20 ng/µL each), Cas9 protein (final concentration 50 ng/µL), and the template sequence (final concentration 5 ng/µL) was microinjected into mouse embryos by the Penn Genetics Transgenic Mouse Core. Born pups were genotyped and sequenced to verify proper insertion of loxP sites. For some experiments (as indicated), mice were fed a high fat diet (Research Diets D12492i: 20% carbohydrate, 60% fat, 20% protein) or high carbohydrate diet (Research Diets D12450Bi: 70% carbohydrate, 10% fat, and 20% protein).

#### 4.1.1. AAV Injections

Mice were injected retro-orbitally at 6–8 weeks of age with adeno-associated virus containing a liver-specific promoter, *thyroxine-binding globulin* (TBG), driving expression of either GFP (AAV-TBG-GFP) or Cre recombinase (AAV-TBG-Cre) (University of Pennsylvania Vector Core) at a dose of 10^11^ genome copies per mouse. For the inducible GKRP-P446L model, *Gckr* floxed mice were injected with 3 x 10^11^ genome copies of either AAV-TBG-hGKRP or AAV-TBG-P446L along with 1 x 10^11^ genome copies of AAV-TBG-Cre. For HKII expression, 1.5 x 10^11^ genome copies of AAV-TBG-HKII were injected along with the 10^11^ genome copies of AAV-TBG-Cre. Diets/experiments were started 2–3 weeks after virus injection.

#### 4.1.2. Glucose tolerance test and Insulin measurements

Mice were fasted for 16h, followed by intra-peritoneal injection of 2 g/kg of 20% glucose. Blood glucose was measured at baseline and 15, 30, 60, and 120 min post-injection via tail bleed using a OneTouch glucose meter and glucose strips. Insulin concentration was measured from the baseline and 15 min samples using an Ultra-Sensitive Mouse Insulin ELISA Kit (Crystal Chem).

#### 4.1.3. Hepatic Glycogen measurement

50 mg of liver was homogenized in 250 µL of 6% perchloric acid in a TissueLyser (Qiagen) at 30 Hz for 3 minutes. The homogenate was centrifuged at 17,000 *g* for 10 min at 4 °C. 125 µL of the supernatant was transferred into a new tube and diluted with 125 µL deionized water. The sample was again centrifuged at 17,000 *g* for 10 minutes at 4 °C and 400 µL was transferred to a new tube. 8.38 µL of 10 N KOH was added to neutralize the pH. The sample was centrifuged at 17,000 *g* for 10 min at 4°C and the supernatant was transferred to a new tube. 20 µL of each sample was treated with 100 µL of amyloglucosidase (Sigma A7420, 1mg/mL) and incubated at 40 °C for 2 hours with constant shaking. For the glucose assay, 10 µL of sample/standard (Sigma G6918) were loaded onto a 96 -well clear plate with 200 µL glucose assay reagent (Sigma G3293). 10 µL of the pre-digested glycogen samples were also loaded to determine background glucose levels. Absorbance was measured at 340 nm and concentration was calculated via standard curve and normalized to liver weight.

#### 4.1.4. Hepatic TAG assay

100 mg of liver was homogenized in 400 µL of cell lysis buffer (50 mM Tris-HCl, 140 mM NaCl, 0.1% Triton-X100, pH 7.4) with a TissueLyser (Qiagen) at 30 Hz for 2 min. Refed samples were diluted 10x while overnight fasted samples were diluted 25x in cell lysis buffer. 20 µL of sample were loaded onto a 96-well clear plate along with a triglyceride standard (Fisher Scientific #23-666-422). 20 µL of 1% sodium deoxycholate was added to each well and the plate was incubated at 37°C for 10 min. 200 µL of Infinity Triglyceride Reagent (Thermo Scientific TR22421) was added and the plate was incubated at 37°C for 10 min. Absorbance was measured at 500 nm and concentration was calculated via standard curve and normalized to liver weight.

#### 4.1.5. Plasma TAG and Cholesterol

Blood samples were collected in heparin coated collection tubes and plasma was separated by centrifuging for 15 min at 5,000 *g* at 4 °C. Plasma samples were diluted 5x with deionized water. 20 µL of sample was loaded into a 96-well clear plate along with either a triglyceride (Fisher Scientific #23-666-422) or cholesterol (Pointe Scientific C7509-STD) standard. For the triglyceride assay, 20 µL of 1% sodium deoxycholate was added to each well and the plate was incubated at 37 °C for 10 min. 200 µL of Infinity Triglyceride Reagent (Thermo Scientific TR22421) or Infinity Total Cholesterol Reagent (Thermo Fisher Scientific TR13421) was added and the plates were incubated at 37 °C for 10 min. Absorbance was measured at 500 nm and concentration was calculated via standard curve.

#### 4.1.6. De novo lipogenesis

Mice were fed a HCD (Research Diets D12450Bi) for 1 week and then fasted overnight (16 hours). Mice were refed HCD for 3 hours and then injected with 20 µL/g body weight deuterated water with 9 mg/mL NaCl before being allowed to feed for an additional 3 hours. Mice were sacrificed and their livers and blood plasma were harvested. Deuterium incorporation into palmitate was measured via mass spectrometry (Metabolism Tracer Resource, University of Pennsylvania Diabetes Research Center). For fasted samples, mice were injected with the deuterated saline 3 hours prior to the completion of the overnight fast and sacrificed once the fast ended.

#### 4.1.7. Cholesterol Synthesis

Mice were fed HCD for 1 week and fasted overnight. Immediately prior to refeeding, mice were injected 20 µL/g body weight deuterated water with 9 mg/mL NaCl and then refed HCD and provided drinking water with 4% deuterated water for 24 hours. Mice were sacrificed and their livers and blood plasma were harvested. Deuterium incorporation into cholesterol was measured via mass spectrometry (Metabolism Tracer Resource, University of Pennsylvania Diabetes Research Center).

### 4.2. AAV preparation

AAV8-Rep/Cap and Ad5-Helper plasmids were obtained from the Stanger Lab [37]. AAV expression plasmids for human *GCKR* and *HK2* were designed and ordered from VectorBuilder along with a vector to express the P446L variant. HEK293T cells were plated in 10 cm dishes and grown to 90-100% confluency. Cells were replenished in 6.5 mL DMEM (Cytiva SH30081) with 2% fetal bovine serum (Sigma-Aldrich F0926), 1% GlutaMAX (Thermo Fisher 35050061), and no antibiotics. To make each virus, 6 µg AAV8-Rep/Cap plasmid, 6 µg Ad5-Helper plasmid, and 6 µg of the particular AAV expression plasmid along with 54 µg of polyethylenimine (Linear, MW 25,000, Polysciences 23966) were mixed in 3.5 mL Opti-MEM (Thermo Fisher 31985070) and vortexed briefly to mix. The plasmid/PEI complex was incubated for 15 minutes at room temperature and then added to the cells dropwise with gentle shaking to mix. After incubation for 6 days, the cells and media were collected and centrifuged at 1600 *g* for 15 minutes. The supernatant was transferred into a fresh tube and benzonase (Sigma-Aldrich# E1014) was added at a 1:40,000 ratio, mixed by inversion, and incubated for 30 minutes at 37 °C. The sample was centrifuged again at 1600 *g* for 15 minutes and the supernatant was filtered through a 0.22 µm vacuum filtration system (Sartorius 180E01). A 40% solution of polyethylene glycol 8000 (Bioworld 41600048) in 2.5 M NaCl was added to the filtered media at a 1:4 ratio and incubated overnight at 4 °C. The solution was centrifuged at 3000 *g* for 15 minutes and the supernatant was carefully removed from the precipitated AAV. The precipitate was homogenized in ∼10 µL PBS per 10 cm dish via pipetting and filtered in a 0.45 µm filter column (Corning 8162). Viral titre was determined via qPCR using AAV-TBG-GFP as a standard.

### 4.3. mRNA isolation, real-time qPCR and RNA sequencing

Total RNA was isolated from frozen liver samples using the RNeasy Plus kit (Qiagen Cat# 74134), including the DNase digestion step (Qiagen Cat #79254). Complementary DNA was generated using M-MuLV <u>reverse transcriptase</u> and relative gene expression was quantified by <u>real time qPCR</u> using the SYBR Green dye-based assay (ABI). Hepatic RNA samples were also processed for bulk RNA sequencing (Novogene). Differential expression was analyzed with the Illumina DESeq2 program [38]. Gene ontology was performed by inputting the lists of significantly up- and downregulated genes (FC ≥ 2 or ≤ 2 and adjusted p value < 0.05) into Enrichr [39–41] and the top 5 GO terms (GO Biological Process 2026) were highlighted with their respective p value.

### 4.4. Protein isolation and western blotting

20-25 mg of liver was homogenized in RIPA buffer (50mM Tris-HCl, 1% Triton x100, 0.5% Sodium deoxycholate, 0.1% SDS, 150 mM NaCl, 2 mM EDTA) supplemented with protease inhibitor cocktail tablets, and phosphatase inhibitor cocktail II and III (Sigma Aldrich #P5726 and #P0044) in a TissueLyser (QIAGEN) at 30 Hz for 2 minutes. Protein concentrations were determined via the Pierce ™ BCA Protein Assay Kit (ThermoFisher). 30 µg of protein was separated using 4-15% Mini-PROTEAN TGX pre-cast gels. Primary antibodies used were HSP90 (Cell Signaling Technology #4874), Gck (Proteintech #19666-1-AP), GKRP (Cell Signaling Technology #14328), and HKII (Cell Signaling Technology #2867).

### 4.5. Quantification and Statistical Analysis

Statistical analysis was performed using One-way ANOVA coupled with Tukey’s multiple comparison test when more than two groups were compared, Two-way ANOVA with Tukey’s multiple comparison test when two conditions were analyzed, and unpaired two-tailed Students’ t-test when only two groups were being compared. All data are presented as mean ± standard error of mean. */† indicates p-value < 0.05, **/†† indicates p-value < 0.01, ***/††† indicates p-value < 0.001, and **** indicates p-value < 0.0001.

## Data Availability

The RNA-seq dataset generated in this study has been deposited in the Gene Expression Omnibus (GEO) under accession number GSE334975. All other data supporting the findings of this study are available from the corresponding author upon request.

## Acknowledgments

This work was supported by NIH grants UM1 DK126194 (P.S. & P.M.T.), K01 DK111715 (P.M.T.), and R01 DK125497 (P.M.T.) as well as funds from the Cox Research Institute. We thank the University of Pennsylvania Diabetes Research Center (DRC) for the use of the Metabolic Tracer Resource (NIH DK019525) to quantify lipid and cholesterol synthesis as well as the Vector Core for GFP and Cre expressing AAVs. We also thank Drs. Benjamin Stanger and Takeshi Katsuda for providing the plasmids and protocol to create adeno-associated viruses.

## CRediT Authorship Contribution Statement

**Dominic Santoleri:** Conceptualization, Data Curation, Formal Analysis, Investigation, Methodology, Writing – Original Draft, Writing – Review & Editing, Validation, Visualization. **Sarah Traynor:** Methodology, Writing – Review & Editing. **Ryan Calhoun:** Data Curation, Visualization, Writing – Review & Editing **Matthew J. Gavin:** Investigation, Writing – Review & Editing. **David Merrick:** Methodology, Resources, Writing – Review & Editing. **Patrick Seale:** Conceptualization, Funding acquisition, Project administration, Resources, Supervision, Writing – Review & Editing. **Paul M. Titchenell**: Conceptualization, Funding Acquisition, Methodology, Project Administration, Resources, Supervision, Writing – Review & Editing

