## Supplemental Figures for "Glucokinase activity suppresses hepatic cholesterol synthesis and triglyceride accumulation: A new model for the effects of the GKRP P466L common human variant"

| Reported Trait | p value | Beta | Odds Ratio | 95% Confidence Interval | Citation |
|---|---|---|---|---|---|
| Triglycerides | $1.5 \times 10^{-9}$ | | | | Orho-Melander et al. 2008 [19] |
| | $5 \times 10^{-4}$ | | 3.69 | 1.61 - 5.81 | Vaxillaire et al. 2008 [11] |
| | $2 \times 10^{-239}$ | 0.115 | | | Willer et al. 2013 [11] |
| Metabolism-associated steatotic liver disease | $1.06 \times 10^{-10}$ | | 1.278 | 1.186 - 1.377 | Anstee et al. 2020 [15] |
| Metabolism-associated steatohepatitis | $3.78 \times 10^{-7}$ | | 1.302 | 1.176 - 1.442 | Anstee et al. 2020 [15] |
| Total Cholesterol | $3 \times 10^{-42}$ | 0.051 | | | Willer et al. 2013 [11] |
| HOMA-IR | $5 \times 10^{-5}$ | | | | Orho-Melander et al. 2008 [19] |
| | $4 \times 10^{-6}$ | | -5.45 | -7.66 - -3.18 | Vaxillaire et al. 2008 [11] |
| Fasting Insulin | $5 \times 10^{-5}$ | | -3.9 | -5.72 - -2.04 | Vaxillaire et al. 2008 [11] |
| Fasting Glucose | $9 \times 10^{-9}$ | | -1.27 | -1.7 - -0.84 | Vaxillaire et al. 2008 [11] |

**Figure S1: Summary of GWAS traits associated with rs1260326, the GKRP P446L variant**

A.

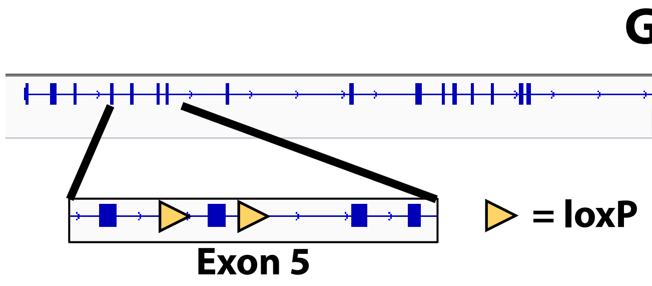


B.

ttctggaattttaatgaatagcttagtcagagaggatagtcaagaggatttctttctccagattactgttctttttgctacctcacct
ggcctatgctcgcctccattttatgatgatgagaagaagaaggtgttccaagtctccctctgacctgggcttctttccatcctca
ggagccagatgggggcctggtggtgctgagtggagggggcacctctggtcgtatggcattccttatgtctgtgagtacctta
attcctggcttctcccttctcctcctggggatagcccccagaacgaaaaggctcatctgaaataaggtctctcaatagatgtat
tctctccccacttcctctccaaaacctggagctcaggatggagtcagatttctctggccccactagagccccggcactctctt
gttaacatcctcacatcttgtgcatcactacgtaatcatttccaagttgcatctttaagtggcggactccaaacaagtataactt
cgtatagcatacattatacgaagttattagggttaagcattggcctaagaagaccattatgcaccttgatcctctaggacctg
agaacccgtgttatgtcctttttaatcactctgtcggcatattccctccccctggacagGTGTCTTTCAACCAGCTG
ATGAAAGGTCTGGGACAAAAACCTCTTTACACATACCTCATTGCAGGGGGTGACAG
gtaagttaccccgacctgtactcgtggcctgaatgttttttgtttgtttgtttgttttttgttttttttaattgtatatgagtatactgtag
ctgtcttcagacataccagaagtgggcattggatcccattacagatgattgtgagccaccatgtggttgctggaaattgaact
cagtacctttggaggaacaatcagtgctcttaactgctgagccatctctccagcccccaaatgtctttttagaggctaaatttg
aatgaatttagtcaagcattatattgaacctgccttagtctccaacatcccataacttcgtatagcatacattatacgaagttatt
tgtggctttgtatctggccctgtcacagatgtttattgtgtgtcacgcctatcacagagcctgttaaacagtaggcgctgttcaat
gatagctttcctccatccctttctgttagtgtacctttcctaaccaccagtaggtatctgctccttacctaataccactgacctcgg
ctgagtttgcacccttaggtctgttgtagcctctcgggaacggacagaagatagcgccctacacggaatcgaggagctga
agaaggtctgtgtctcctgttactacttacctggagatgcccatatattcctttacatagggttagaaaacccccaaccttcag
ccaaggcctggctcctgctctggtcctgattttctcacacactagatttgtaccctggagaccctaatgcacccctttcattccta
ggtggctgctgggaaaaagagagtggtcgttataggcatttccgtgggactctctgtaagtgtgaggtcaggaggggtg

Homology sequence

gRNA site #1: ggcggactccaaacaagtta

gRNA site #2: ttagtctccaacatcccttg

LoxP sequence: ataacttcgtatagcatacattatacgaagttat

*Gckr* Exon 5

**Figure S2: Design of *Gckr* floxed mice:**
**A)** Schematic of the mouse *Gckr* gene with exon 5 enlarged. Exon 5 was removed via CRISPR and replaced with a template containing loxP sites flanking it (detailed as triangles). **B)** Sequence for anti-sense template to create the floxed gene. The loxP sites were placed within the gRNA sites to prevent further Cas9 cutting once the template was incorporated into the genome.

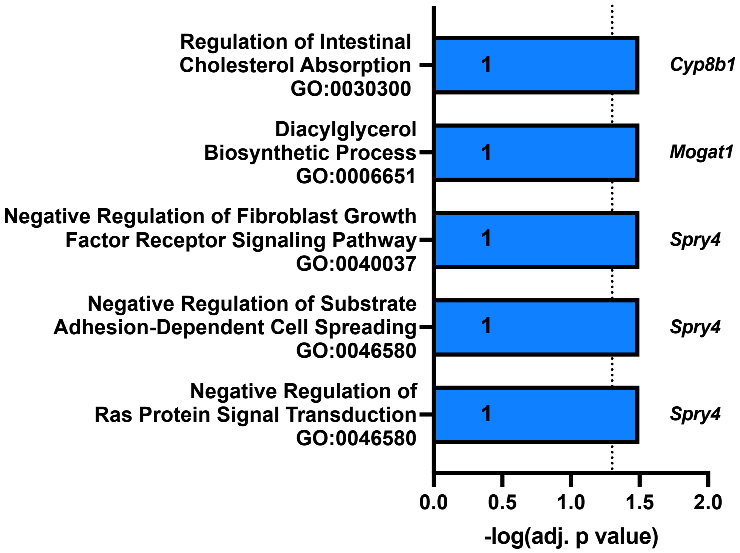


**Figure S3: Top 5 downregulated gene ontology pathways from RNA-seq dataset (GO Biologic Process 2026).** The number within each bar represents how many genes within each pathway were significantly down in the RNA-seq dataset. The list of those genes is presented next to each bar.
